# On statistical modeling of sequencing noise in high depth data to assess tumor evolution

**DOI:** 10.1101/128587

**Authors:** Raul Rabadan, Gyan Bhanot, Sonia Marsilio, Nicholas Chiorazzi, Laura Pasqualucci, Hossein Khiabanian

## Abstract

One cause of cancer mortality is tumor evolution to therapy-resistant disease. First line therapy often targets the dominant clone, and drug resistance can emerges from preexisting clones that gain fitness through therapy-induced natural selection. Such mutations may be identified using targeted sequencing assays by analysis of noise in high-depth data. Here, we develop a comprehensive, unbiased model for sequencing error background. We find that noise in sufficiently deep DNA sequencing data can be approximated by aggregating negative binomial distributions. Mutations with frequencies above noise may have prognostic value. We evaluate our model with simulated exponentially expanded populations as well as data from cell line and patient sample dilution experiments, demonstrating its utility in prognosticating tumor progression. Our results may have the potential to identify significant mutations that can cause recurrence. These results are relevant in the pretreatment clinical setting to determine appropriate therapy and prepare for potential recurrence pretreatment.

## 1 Introduction

Every extant organism is the result of over three billion years of evolution. Complex organisms consist of cells whose functions are regulated by a large number of interconnected pathways that ensure cellular, tissue, and organ homeostasis. Cancer is a result of the breakdown of this process in a single cell, which results in its unregulated growth. In most cases, the immune system is able to detect and eliminate such aberrant cells. Sometimes, however, a clone escapes this surveillance and manifests as clinically detectable disease ^40^. Consequently, most clinically diagnosable tumors are clonal, i.e. they grow clonally from a single cell that finds a path to circumvent the body’s defense mechanisms. The growing tumor accumulates mutations, most of which have low fitness and therefore are found at low frequencies, outcompeted by the dominant clone.

The clonal expansion process, which underlies genomic diversification within a tumor, was first studied by Salvador Luria and Max Delbrück. They designed a simple system of single-cell organisms to investigate patterns of mutation accumulation. Their rigorous quantitative methodology led them to discover that mutations arise randomly and their numbers follow a distinct probability distribution^24^. As the cell population in the tumor diversifies, it is able to explore the fitness landscape. Studying the dynamics of this genomic heterogeneity can yield insight into when the clonal expansion started, how fast the population evolved, and whether specific genomic alterations were selected in a particular host or under a treatment regimen.

The principal biochemical mechanisms in cancer are often recurrent across tumors in different tissues. For example, aberrations leading to unregulated cell growth or inactivation of the apoptotic pathway (cell suicide) are common to almost all tumors. Given the limits within which cells are regulated, the growing tumor has access to only a finite number of pathways that it can alter. As a result, tumors arising from different cells of origin often harbor identical genetic mutations, which alter the same pathways, and often have similar prognostic consequences ^5^.

First line therapy drugs target a tumor’s dominant, fastest growing clone. Drug resistance often emerges from the rise of preexisting clones that harbor potential driver mutations that gain evolutionary fitness via therapy-induced natural selection. It has been shown that the presence of drug-resistant sub-clones in the primary tumor prior to therapy may be a strong predictor of poor survival, with direct implications for disease management ^28,34,37,44^. As cancer therapy moves towards individualized treatment, it is important to identify and understand the role of such mutations, some of which may have prognostic value. Such potentially prognostic mutations are commonly identified using targeted deep sequencing of the tumor DNA in clinical settings, and their sensitive detection relies on the accurate analysis of background noise, specifically DNA sequencing errors.

Studying the evolution of chronic lymphocytic leukemia (CLL) under therapy is an illuminating example of these approaches ^19,20^. CLL is the most common leukemia in adults and its clinical course ranges from asymptomatic disease that never requires therapy to rapidly progressive disease that requires intensive treatment. Genomic alterations in CLL follow a time ordered process ^45^. Patients who harbor genomic defects in the *TP53* gene, which regulates many pathways including the cell suicide or apoptotic pathway, are considered at high risk of failing conventional therapies ^35^. Such patients are good candidates for stem cell transplant or new gene-specific therapeutics ^2,39^. The presence of such secondary mutations in genes such as *TP53* is often assessed using traditional Sanger sequencing that only provides sufficient power to detect mutations present in at least 20% of leukemia cells ^32^. To assess the presence of *TP53* prognostic mutations at lower abundances in newly diagnosed CLL patients, we used deep sequencing and evaluated thousands of leukemia cells and identified small *TP53* mutations that were missed by traditional methods such as Sanger sequencing ^34^. We found that *TP53* mutated sub-clones identified before treatment became the predominant population at the time of CLL relapse, as a result of therapy induced selection pressure. These results suggest that tumors harboring small *TP53* mutations have the same clinical phenotype and risk of failing therapy as those with *TP53* defects in the dominant clone ^27,34^, and their early detection is essential for the identification and management of high-risk CLL patients ^11^.

These results are also pertinent to other hematological malignancies where the presence of leukemia-associated mutations in remission is associated with significantly increased risk of relapse and poor survival^31,37^. These data lead to the conclusion that it is imperative to identify alterations that induce therapeutic resistance in leukemia patients in the early stages of disease in order to properly guide individualized therapy with the goal of preventing disease relapse. However, the detection of mutations at low allele frequencies (e.g., 1 mutation in 10,000 cells) is hindered by the lack of a precise model of noise in diagnostic sequencing assays.

Targeted sequencing is the most commonly used method to track prognostic markers in both clinical and basic research applications ^10^. However, finding such mutations in sequencing reads is often confounded by misreading a base in the sequencing instrument or mis-incorporation of DNA bases (nucleotides) during library enrichment by polymerase chain reaction (PCR) amplification cycles. More accurate sequencing protocols, which perform overlapping reads of the same genomic DNA region, allows the merging of such reads for improved accuracy. This facilitates correcting errors accumulated in the sequencer, while leaving uncorrected PCR errors that arise during library preparation steps ^4,46^.

The challenge in identifying potentially functional sub-dominant mutations is to determine the sensitivity thresholds of sequencing platforms, i.e. the depths above which PCR errors happen with a probability below a statistical cut-off. Such thresholds can be estimated by hypothesizing that all variants are due to errors and using deviations from this null hypothesis to indicate the presence of true variants. This can sometimes be confounded by the fact that different sequencing errors occur at different rates ^3,6^. Hence a single threshold cannot comprehensively test the significance of all variants. As a result, more sophisticated statistical modeling of the background error distribution is necessary.

To model background error one may use different types of error distributions: (i) a single or a linear combination of Luria-Delbrück distributions, characterizing the expected number of spontaneous mutations during tumor growth, where the PCR error rate is assumed to be constant ^17^; (ii) the negative binomial distribution, describing the depth distribution of clones after PCR amplification through a Poisson-Gamma mixture model ^29^; and iii) the beta-binomial distribution, suitable for Bayesian models, where error rates are assumed to follow the Beta distribution ^21^. Although the Luria-Delbruück distribution is expected to better describe the long tail of the error depths, empirical analysis has shown that the negative binomial distribution gives the best fit to the observed error depths based on goodness-of-fit log-likelihood ^34^. The beta-binomial distribution, in conjunction with multiple filtering criteria based on normal control DNA samples, has also been proposed for somatic mutation detection from cancer genomes ^8,9,23,36^.

In this manuscript, we revisit this problem and provide a comprehensive model that illustrates how aggregate negative binomial distributions describe PCR error depths in ultra-deep targeted sequencing. We test our model with *in silico* as well as cell line and patient dilution experiments, and propose a highly sensitive, mutation-specific approach to detect true mutations, without the need for control data from un-mutated (wild type) normal tissue DNA.

## 2 Methods

### Derivation of the error depth distribution

Here we will only be discussing the distribution of low frequency errors in deep DNA sequencing analysis of tumor samples. Let us assume an experiment in which S independent tissue samples are subjected to ultra-deep sequencing. DNA sequencing of tumor samples produces strings of nucleotides (A, C, G, and T) of 100-200 base-pair length that correspond to the DNA sequences of different sections of the genome in the tumor sample. These sequences of DNA reads are mapped to a “reference” genome and deviations/mismatches are identified as potential mutations. Ideally, the reference sequence is the sequence from the patient’s “germ-line”, usually obtained from blood or some other tissue with normal cells. The sequencing read depth is the average number of reads that map to the same locus (section of the genome). At a nucleotide, three potential single base substitutions can occur: A (adenine) → C, G, T, or C (cytosine) → A, G, T, or G (guanine) → A, C, T, or T (thymine) → A, C, G. Alternately, there might be an insertion (addition of one or more A, C, G, T nucleotides) or a deletion (loss of A, C, G, T nucleotides). All of these will henceforth be referred to as variants. We want to derive the posterior probability distribution for these variants, assuming they are stochastic, i.e. they represent noise (statistical random errors).

Suppose that, at a genomic DNA locus, we see *n_i_* such variant reads amongst *N_i_* total reads. The distribution of *n_i_* follows a binomial distribution, Bino(*n_i_|N_i_, θ*), where *θ* is the *a priori* probability of a variant’s occurrence. Let 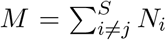 be the total number of reads across samples at that locus and 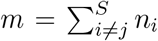 be the total number of variant (erroneous) reads across samples at that DNA locus. Then, the posterior predictive *p* value for having detected a true mutation in sample *j*, given *S* – 1 other samples, can be obtained from the posterior probability distribution:

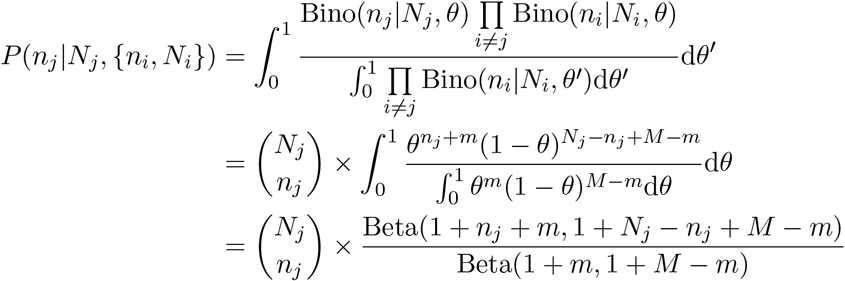

where Beta indicates the Beta function. Simplifying the algebra yields the beta-binomial distribution,

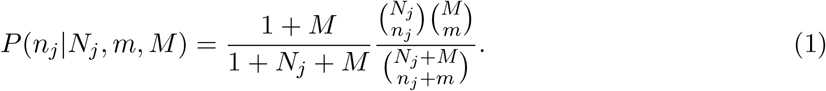

Variations of equation (1) have been previously derived for sequencing depths > 100× ^8,9,36^. Today, it is possible to do ultra-deep sequencing, where *N_i_* > 5,000×. In such cases, for low frequency variants, we can assume that *n_i_ ≪ N_i_*. Therefore, we can use Stirling’s approximation, and estimate 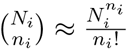. Equation (1) can then be approximated by

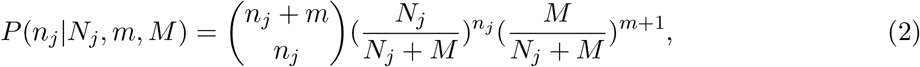

which equals 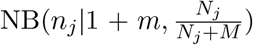, with NB indicating the negative binomial distribution, and where 1 + *m* and 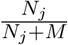 are its two parameters, which we can interpret as the number of detected errors and the *a priori* probability of success in detecting an error, respectively.

### Exponential expansions at varying error rates

An exponentially expanded population is generated through *c* PCR amplification cycles, where each cycle doubles the DNA population. If errors accumulate independently at a rate of *μ* substitutions per site per cycle, the average error depth (i.e. the average number of reads harboring errors) is 2*^c^μ*. For *S* such populations, the error depth distribution is described by equation (1), or is approximated by a negative binomial distribution, 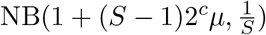, as derived above in equation (2).

It is well known that different types of PCR errors occur at different rates. For example, transitions, that exchange two-ring purines (A and G) or one-ring pyrimidines (C and T) are more common than transversions, which replace an A or G with one of C or T. Assuming *R* independent rates, the observed number of variants *D(v)*, with error depth *v* is then given by,

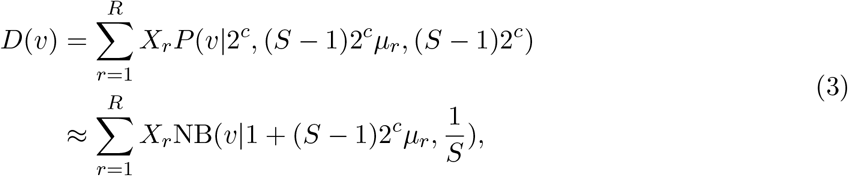

where *X_r_* represents the number of variants that occur with rate *μ_r_*. Since error rates are often unknown and sequence context dependent, we can alternatively bin the variants based on their average error depth across samples and write *D(v)* as

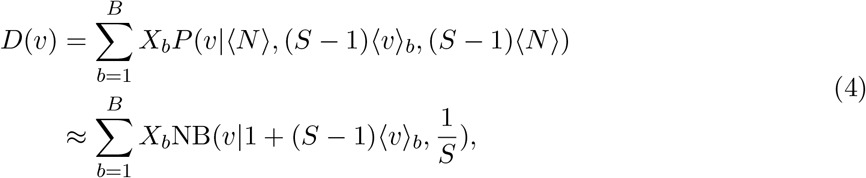

where *B* is the number of bins, *X_b_* is the number of variants in each bin, and ⟨*N*⟩ is the average sequencing depth across *S* samples. It has been shown that the sum of negative binomial distributions with equal success probabilities is also a negative binomial distribution, though with a random parameter ^7,43^. Thus, the approximation of *D(v)* in equations (3) and (4) with sums of negative binomial distributions that have success probability of 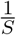, suggests empirical observations ^34^

## 3 Data

In the first experiment, a series of dilutions was generated using the SU-DHL-6 cell line (Diffuse Large B-Cell Lymphoma), which carries a heterozygous (one allele altered) TP53-Y234C missense mutation (one that changes an amino acid in a protein sequence) ^26^. The cells were serially diluted at (1:10, 1:10^2^, 1:10^3^, 5:10^4^, 1:10^4^, 5:10^5^, and 1:10^5^) by mixing the cell line DNA with *TP53* wild-type genomic DNA from a healthy donor. The *TP53* mutation locus was sequenced at depths of 10,000× (10K×), 100,000× (100K×), and 1,000,000× (1M×).

In the second experiment, genomic samples from 18 healthy individuals as well as samples from undiluted and 1:10^3^ diluted cancer cells from a CLL patient, harboring a heterozygous *SF3B1*-K700E missense transition substitution were analyzed and the *SF3B1* mutated locus was sequenced at a mean depth of 620,000 ×.

For both experiments, each cell line dilution and patient sample was barcoded and targeted with amplicon multiplexed sequencing using the Illumina MiSeq (2 × 150 bp) (Genewiz, South Plainfield, NJ). The primers were designed so that the paired-end reads substantially overlapped with each other and each read pair was merged to correct sequencing errors. The merged reads were mapped to the human reference genome (hg19) using the Burrows-Wheeler Aligner (BWA) alignment tool^22^, and all variable sites were identified using an inclusive variant caller, adapted from the SAVI algorithm^41^.

## 4 Results

### Simulated data

We generated a set of *in silico* experiments with exponentially expanded populations starting from a single, homogenous, 100 base-long sequence of binary bases. Each population was aggregated from four expansions that followed error rates of 10^-3^, 10^-4^, 10^-5^, and 10^-6^ substitutions per site per cycle. The number 12, 14, and 18 of cycles were chosen to produce populations with 16,384, 65,536 and 1,048,576 total reads respectively. Each experiment contained 50 independent populations (*S* = 50) and for each experiment, *D(v)*, the expected number of variants with depth v was calculated using equations (3). This experiment was repeated 100 times. Figure 1 shows the results, as well as statistically significant *χ*^2^ *p* values indicating high accuracy of the estimates from both the beta-binomial model and its NB approximation.

**Figure 1:**
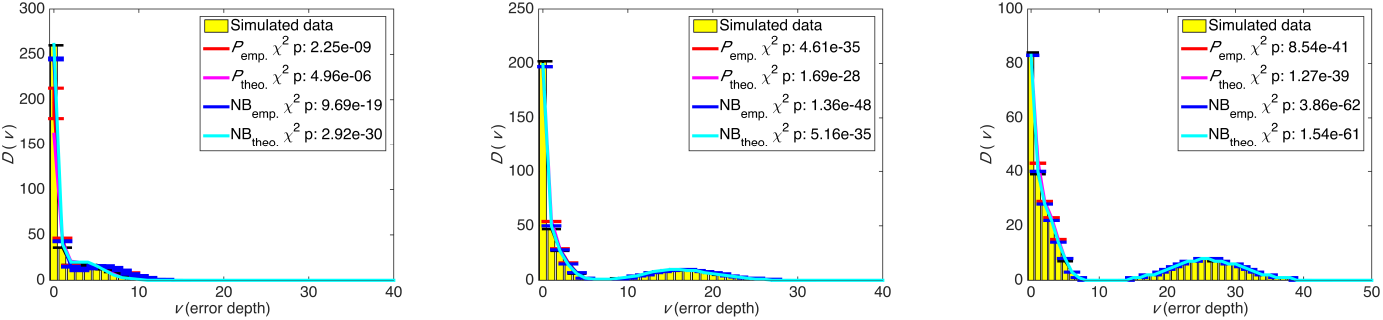
Number of variants with error depth of *v* from aggregated simulated cycles of PCR amplification at four error rates: 12 cycles (left), 14 cycles (middle), and 18 cycles (right). *P*_theo_. and NB_theo_. are calculated using equation (3), and *P*_emp_. and NB_emp_. are calculates using equation (4). The *χ^2^* test was used to compare the distributions.

### Dilution experiments

We removed the real diluted *TP53* mutation from cell line sequencing data, and arranged the erroneous variants based on their depth in 5×-sized bins. We then counted the number of variants *X_b_* in each bin, and calculated *D(v)* using equation (4). Figures 2, 3, and 4 show the results for sequencing depths of 10K×, 100K×, and 1M×, indicating statistically significant *χ*^2^ *p* values that show a strong concordance between estimates from the beta-binomial model, its NB approximation, and ultra-deep sequencing data. Distinguishing transitions and transversions further clarified the importance of classifying variants using sequencing depth as a proxy for the error rates. We obtain similar results from modeling the ultra-deep sequencing data from the *SF3B1* locus (Figure 5).

**Figure 2:**
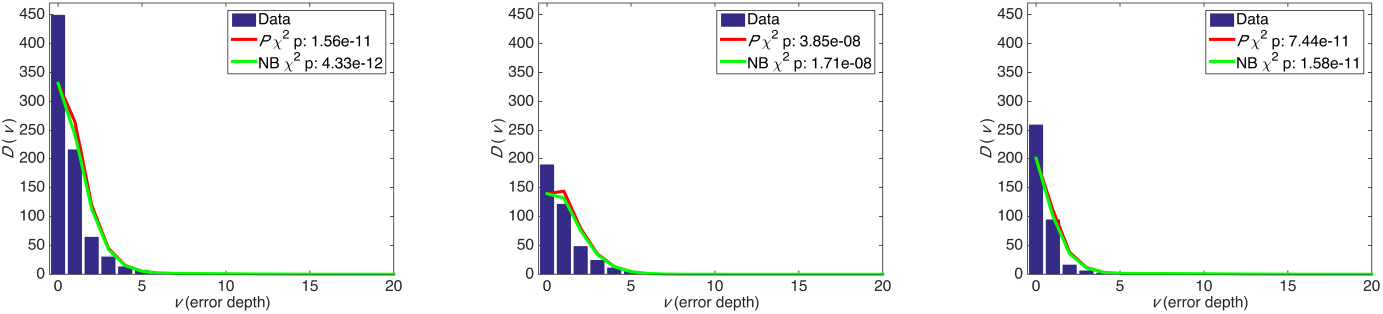
Error depth distribution in ultra-deep sequencing of a *TP53* locus at 10,000× for all variants (left), transitions (middle), and transversions (right).

**Figure 3:**
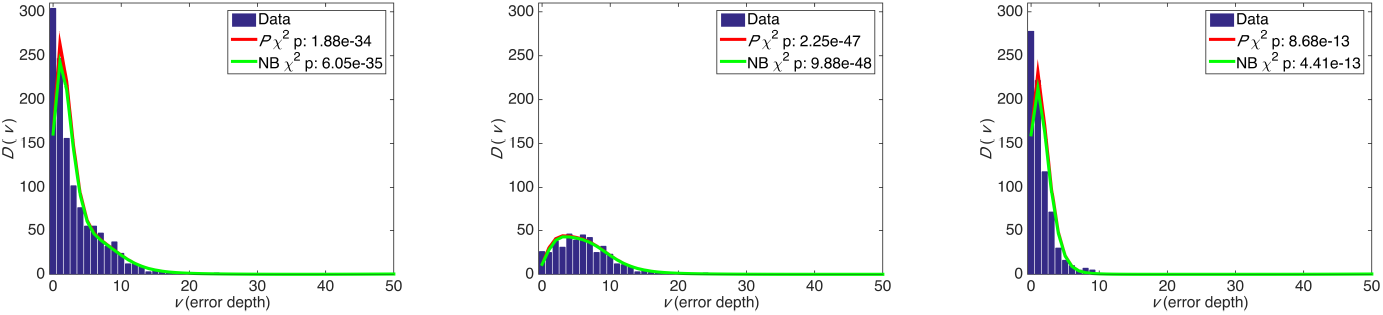
Error depth distribution in ultra-deep sequencing of a *TP53* locus at 100,000× for all variants (left), transitions (middle), and transversions (right).

**Figure 4:**
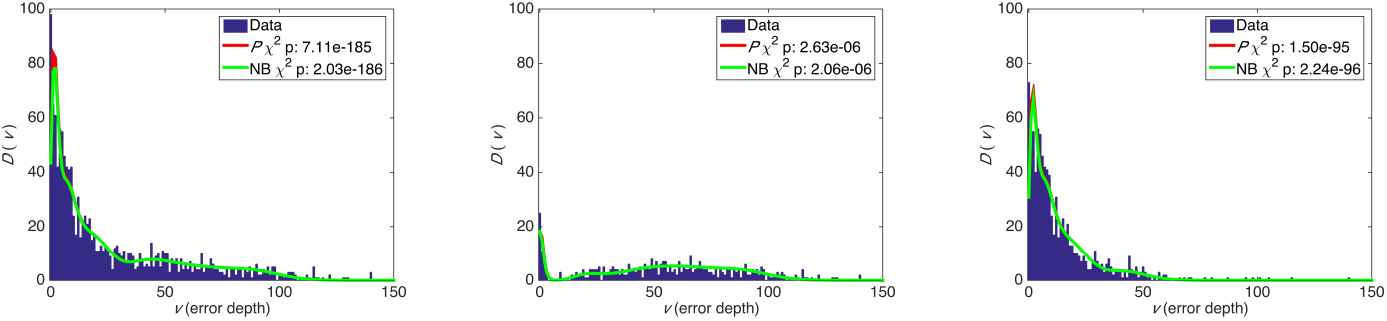
Error depth distribution in ultra-deep sequencing of a *TP53* locus at 1,000,000× for all variants (left), transitions (middle), and transversions (right).

**Figure 5:**
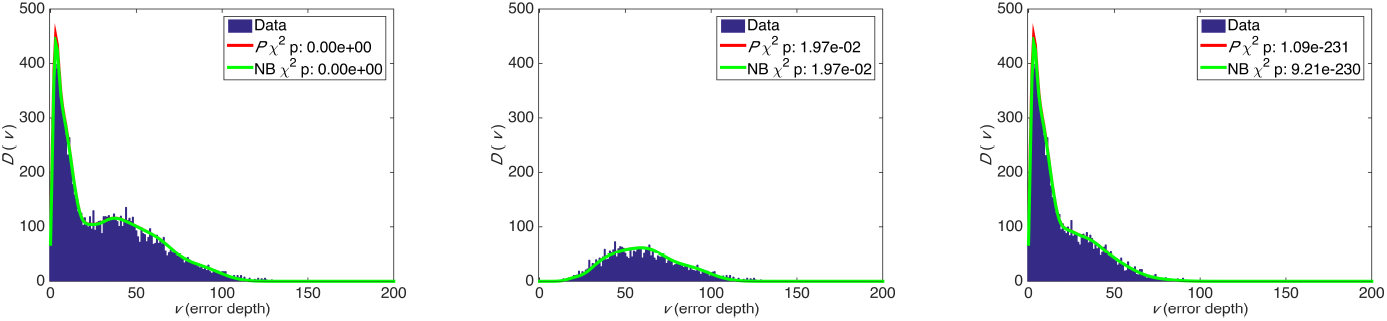
Error depth distribution in ultra-deep sequencing of a *SF3B1* locus at mean 620,000× for all variants (left), transitions (middle), and transversions (right).

### Detecting true mutations

We propose two comprehensive approaches to assess the presence of true mutations at very low abundance relative to background. Our methodology does not require matched normal samples or extensive filtering based on variant annotation resources.

First, having established an accurate model to describe the sequencing error distribution, a threshold is determined above which sequencing errors happen with a probability below an established statistical cut-off. These thresholds can be derived from all variants or a subset of variants, for example, only transitions or transversions. Figure 6 shows such thresholds for detecting the *TP53-*Y234C transition mutation in dilution experiments, where we are able to identify the mutation in abundances as low as 5:10^4^ at 10K× and 100K×, and 1:10^4^ at 1M×, without any false positive calls. As shown in Figures 2, 3, and 4, there is better sensitivity for detecting a transversion substitution.

**Figure 6:**
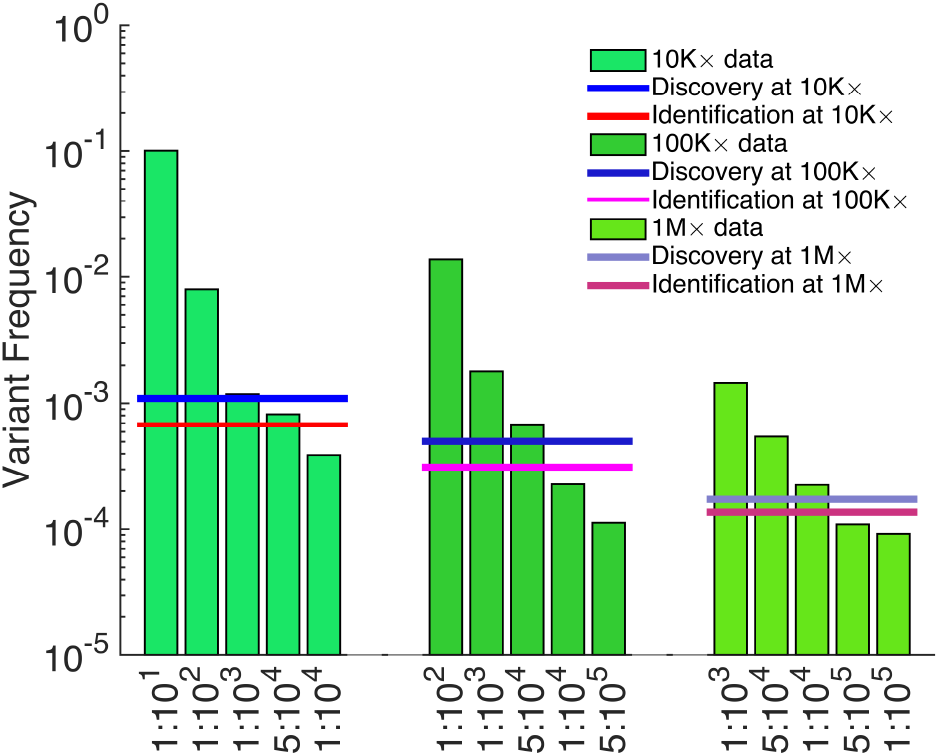
Sensitivity of detecting *TP53*-Y234C mutation dilutions. Assessing the presence of a variant requires correcting for multiple hypotheses based on the number of sequenced genomic positions (Bonferroni correction). Testing the presence of a discovered variant does not require such a correction; here, the *p* value of significance is set at 0.01.

In the absence of matched normal samples, this approach is especially practical for identifying mutations that may exist in more than one tumor sample. Its application to 309 newly diagnosed CLL patients identified small sub-clonal prognostic mutations in four frequently mutated drivers of this neoplasm, present in 2 out of 1,000 wild-type alleles. These mutations were missed by traditional Sanger sequencing, but were validated by independent deep sequencing and allele-specific PCR ^33,34^.

Second, we tested an individual mutation in each sample against all other sequenced samples and calculated the cumulative *P* using equation (1). After correcting for multiple hypotheses using the Benjamini and Hochberg method ^1^, we generated a list of variants that satisfied a pre-determined false discovery rate. This approach is particularly powerful in identifying patient-specific mutations. We assess the presence of the *SF3B1*-K700E mutation in patient samples, and find the probability of observing the mutation in 1:10^3^ CLL dilution to be extremely significant compared to controls (Table 1). This approach can accurately identify sample-specific mutations by comparing multiple samples at the same exact mutated base.

**Table 1:** Presence of the *SF3B1-K700E* mutation in undiluted and diluted patient samples are tested against 18 samples that harbor wild-type allele.

| Sample | Variant Depth ($v$) | Total Depth | Variant Frequency | Cumulative $P$ | FDR | Cumulative NB |
| --- | --- | --- | --- | --- | --- | --- |
| Control | 64 | 711703 | 0.00009 | 9.48E-01 | 9.48E-01 | 8.66E-01 |
| Control | 62 | 642586 | 0.00010 | 8.45E-01 | 9.48E-01 | 6.85E-01 |
| Control | 74 | 717154 | 0.00010 | 7.02E-01 | 9.37E-01 | 5.09E-01 |
| Control | 94 | 630510 | 0.00015 | 2.95E-03 | 9.43E-03 | 5.00E-04 |
| Control | 56 | 505857 | 0.00011 | 4.68E-01 | 7.89E-01 | 2.60E-01 |
| Control | 61 | 509147 | 0.00012 | 2.49E-01 | 5.69E-01 | 1.07E-01 |
| Control | 88 | 699082 | 0.00013 | 1.12E-01 | 2.98E-01 | 4.12E-02 |
| Control | 75 | 749932 | 0.00010 | 7.91E-01 | 9.48E-01 | 6.22E-01 |
| Control | 62 | 657036 | 0.00009 | 8.84E-01 | 9.48E-01 | 7.47E-01 |
| Control | 56 | 581178 | 0.00010 | 8.34E-01 | 9.48E-01 | 6.63E-01 |
| Control | 81 | 731934 | 0.00011 | 4.75E-01 | 7.89E-01 | 2.85E-01 |
| Control | 70 | 636485 | 0.00011 | 4.93E-01 | 7.89E-01 | 2.89E-01 |
| Control | 40 | 452271 | 0.00009 | 9.15E-01 | 9.48E-01 | 7.92E-01 |
| Control | 59 | 511932 | 0.00012 | 3.51E-01 | 7.03E-01 | 1.72E-01 |
| Control | 46 | 518211 | 0.00009 | 9.27E-01 | 9.48E-01 | 8.15E-01 |
| Control | 80 | 714670 | 0.00011 | 4.35E-01 | 7.89E-01 | 2.50E-01 |
| Control | 85 | 736865 | 0.00012 | 3.33E-01 | 7.03E-01 | 1.75E-01 |
| Control | 74 | 691495 | 0.00011 | 5.87E-01 | 8.53E-01 | 3.82E-01 |
| CLL | 281058 | 630750 | 0.44559 | 0.00E+00 | 0.00E+00 | 0.00E+00 |
| CLL 1:1000 dilution | 2678 | 440301 | 0.00608 | 0.00E+00 | 0.00E+00 | 0.00E+00 |

In comparison of our method to other published variant calling algorithms, one comparable unbiased method is EBCall, whose implementation is based on beta-binomial distributions and establishing priors from normal sequencing data ^36^. EBCall requires normal samples; therefore, we removed the reads harboring the diluted mutations in the EBCall analysis to simulate matched normal data. EBCall, with a sensitivity-adjusted configuration, successfully identified the *SF3B1*-K700E mutation in 1:10^3^ CLL dilution sample, as well as the *TP53*-Y234C mutation in the least diluted samples at all sequencing depths (i.e. 1:10 in 10K×, 1:10^2^ in 100K×, and 1:10^3^ in 1M×). However, it failed to detect the mutation at higher dilution levels, and also resulted in four false positive calls at 1M×.

## 5 Conclusion

Therapeutic resistance, one of the main causes of eventual disease relapse and mortality in cancer patients, is often associated with natural selection of preexisting resistant clones under treatment ^12,34^. The detection of such low frequency sub-clones is hindered by a lack of precision-tested diagnostic assays.

Allele-specific, real-time PCR assays have been proposed to identify prognostic variants ^15,25,42^. These approaches only target known mutations, and their adaptation to situations with large numbers of variants requires extensive primer calibration. In contrast, high-throughput sequencing provides an unbiased view of tumor heterogeneity and its genomic profile. Various techniques based on unique molecular identifiers have been proposed to correct both polymerase and sequencing errors ^14,16,18,30^ that facilitate distinguishing real mutations from mistakes that arise during amplification. However, the main hurdle in clinical utilization of these approached is the requirement for generating very large numbers of sequencing reads to assemble the genome of a single DNA molecule with high confidence at depth > 2,000×.

Here, we addressed this important problem in cancer therapy by introducing a highly sensitive method to model sequencing noise, which allows the detection of prognostic markers of disease recurrence using ultra-deep targeted sequencing. Our approach is based on interrogating data from multiple tumor samples at identical genomic regions and provides an accurate assessment of the error rate at a given position without relying on normal samples. Instead of establishing a fixed detection threshold for all variants, we directly calculate mutation-specific sensitivities. Overall, since ultra-deep sequencing methods are now routinely implemented in the clinic, we believe that the application of our comprehensive model to tumor samples will increase the speed with which patients can be evaluated during disease surveillance. Our method opens up the possibility of exploring the dynamics of cancer clones after treatment, timing the rise of resistance to therapy, and determining the clinical importance of minimal residual disease assessed from liquid biopsy samples for precise disease management ^13,38^.

## Acknowledgments

The authors gratefully acknowledge the constructive feedback of Mohammad Hadigol and Alexandra Jacunski. R.R. acknowledges funding from the NIH (U54CA193313, R01CA185486, and R01CA179044). H.K. acknowledges support from the ACS (IRG-15-168-01), Rutgers Cancer Institute (P30CA072720), and Rutgers Office of Advanced Research Computing (NIH 1S10OD012346-01A1).

